# Prior Pro-inflammatory Polarization Changes the Macrophage Response to IL-4

**DOI:** 10.1101/2021.04.06.438673

**Authors:** Erin M. O’Brien, Kara L. Spiller

## Abstract

Tissue repair is largely regulated by diverse macrophage populations whose functions are timing- and context-dependent. The early phase of healing is dominated by pro-inflammatory macrophages, also known as M1, followed by the emergence of a distinct and diverse population that is collectively referred to as M2. The extent of the diversity of the M2 population is unknown. M2 macrophages may originate directly from circulating monocytes or from phenotypic switching of pre-existing M1 macrophages within the site of injury. The differences between these groups have not been investigated, but have major implications for understanding and treating pathologies characterized by deficient M2 activation, such as chronic wounds, which also exhibit diminished M1 macrophage behavior. This study investigated the influence of prior M1 activation on human macrophage polarization to an M2 phenotype in response to IL-4 treatment in vitro. Compared to unactivated (M0) macrophages, M1 macrophages upregulated several receptors that promote the M2 phenotype, including the primary receptor for IL-4. M1 activation also changed the macrophage response to treatment with IL-4, generating an M2-like phenotype with a distinct gene and protein expression signature compared to M2 macrophages prepared directly from M0 macrophages. Functionally, compared to M0-derived M2 macrophages, M1-derived M2 macrophages demonstrated increased migratory response to SDF-1α, and conditioned media from these macrophages promoted increased recruitment of endothelial cells in transwell assays. Together, these findings indicate the importance of prior M1 activation in regulating subsequent M2 behavior, and suggest that augmentation of M1 behavior may be a therapeutic target in dysfunctional tissue repair.

**Summary sentence:** M1 macrophages that are switched to an M2 phenotype exhibit a distinct functional phenotype compared to M2 macrophages derived directly from unactivated (M0) macrophages.

## Introduction

Macrophages are key players in the tissue repair process that regulate the growth of new blood vessels, or angiogenesis. Due to their highly plastic nature, macrophage phenotype varies with the shifting phases of healing by changing in response to external cues. The early inflammatory response to injury is dominated by a pro-inflammatory population commonly referred to as “M1,” which then give way to a very different population known as “M2” in later reparative stages. Studies have shown that this phenotype-switching behavior is characteristic of normal tissue repair, and may be particularly important for angiogenesis (1-3). The late stage, reparative macrophages seen in vivo can emerge in several ways: the recruitment and differentiation of newly arriving non-classical monocytes, the phenotypic switching of pre-existing pro-inflammatory macrophages, and the proliferation of these populations (1, 4-9). It is yet unknown to what extent each mechanism is responsible for generating the collective “M2” population, and it is probable that all are required for normal healing. Nonetheless, studies have shown that when macrophages are depleted in the early inflammatory stage, late-stage M2 macrophages are also reduced and healing is inhibited (6, 10). Furthermore, it is known that in several diseases in which healing is stalled, such as in chronic wounds, pro-inflammatory M1-like macrophages are insufficiently activated and also fail to switch to an M2 phenotype (11-16). Thus, M1-to-M2 phenotypic switching is likely a key mechanism in proper healing, and M1 activation may be an important regulator of subsequent M2 polarization. Despite the significance of the M1-to-M2 transition, the effects of M1 activation on subsequent M2 properties and the specific functions of this M1-derived M2 population have not yet been investigated.

It is important to note that macrophage phenotypes can be categorized using a number of nomenclatures, none of which are universally employed or accepted. One approach is to describe phenotypes based on function, such as “pro-inflammatory,” “pro-angiogenic,” “anti-inflammatory,” “pro-healing,” etc. However, this nomenclature can be problematic due to the influence of timing and context on macrophage behavior; for example, M1 macrophages have been described as both anti- and pro-angiogenic (17, 18) and M2 macrophages, which are commonly called anti-inflammatory, propagate type 2 inflammation (19, 20). Alternatively, the dichotomous system of classically activated “M1” and alternatively activated “M2” was originally coined in reference to the polarizing cytokines secreted by Th1 and Th2 cells (21). This M1/M2 system has since expanded to recognize distinct M2 subtypes, including “M2a” (IL-4-activated macrophages), and is especially useful for describing macrophages activated in vitro, though it falls short in describing the true complexity of macrophage phenotypes. Macrophages in vivo can rarely be described as entirely M1 or M2, instead existing somewhere on a wide spectrum of polarization, but the expression of previously-defined M1 and M2 markers determined from in vitro studies can help to identify “M1-like” or “M2-like” features of more complex phenotypes. In this study we activated macrophages in vitro using defined chemical stimuli, and will therefore use the “M1” and “M2” terminologies for these experiments. For more extensive reviews of the pros and cons of various macrophage nomenclature systems, see Murray et al. (22) and Spiller and Koh (23).

Many studies have been conducted to elucidate the roles of M1-like and M2-like macrophages during angiogenesis and healing in vivo. In the early stages of healing, pro-inflammatory circulating monocytes are recruited to the site of injury where they differentiate to M1-like macrophages (1, 24, 25). These early-arriving macrophages have been shown to be crucial for wound vascularization and complete healing (4, 10, 26). After this stage, M2-like macrophages begin to dominate the milieu, further promoting tissue regeneration (10, 27-29). These M2 macrophages are often modeled in vitro via activation with IL-4 (30-34), although other relevant M2-promoting stimuli include IL10 and apoptotic cells (18, 35). Although the specific roles of each phenotype are still being investigated, the current literature suggests that multiple phenotypes work together in a complementary manner to orchestrate normal healing. Indeed, drug delivery systems designed to sequentially release M1-promoting followed by M2-promoting signals have shown promise in promoting angiogenesis and healing (36).

We hypothesized that M1-to-M2 switching generates a unique phenotype that is important for late-stage healing. We first investigated the potential for unactivated (M0) and M1 macrophages to switch to the M2 phenotype in response to IL-4 in vitro. Then, to understand the impact of M1 activation on subsequent M2 behavior, we compared the gene expression profiles of IL-4-induced M2 macrophages derived from M0 or M1 macrophages (called M0→M2 and M1→M2), then conducted several functional assays of macrophage behavior. The results identify M1→M2 macrophages as a distinct phenotype that upregulates select M2 markers and several genes and proteins related to angiogenesis, and exhibits enhanced pro-healing functions.

## Materials and methods

### Monocyte culture and differentiation into polarized macrophages

Primary human monocytes were purchased from the University of Pennsylvania Human Immunology Core, or isolated as previously described (37, 38) from peripheral blood purchased from the New York Blood Center. Monocytes were cultured on ultra-low attachment plates in RPMI-1640 supplemented with 10% human serum and 1% penicillin/streptomycin. Media was additionally supplemented with 20 ng/mL of MCSF (Peprotech, Rocky Hill, NJ, U.S.) to induce differentiation into macrophages. On day 5, macrophages were either maintained in an unactivated M0 state or they were M1-activated with 100 ng/mL of IFNg (Peprotech) and 100 ng/mL of LPS (MilliporeSigma, Burlington, MA, U.S.) for 24-48 hours (**Figure 1A**). For IL-4Rα flow cytometry, an additional M2 group, activated with 40 ng/mL of IL-4 (Peprotech) and 20 ng/mL of IL-13 (Peprotech) was included. For dose response experiments, M0 and M1 macrophages were treated with four doses of IL-4 (10 ng/mL, 1 ng/mL, 0.1 ng/mL, and 0.01 ng/mL) for six hours. To induce the M0→M2 and M1→M2 phenotypes, M0 and M1 macrophages were treated with 10 ng/mL of IL-4 (Peprotech) on day 6, respectively, for an additional 24-72 hours.

**Figure 1.**
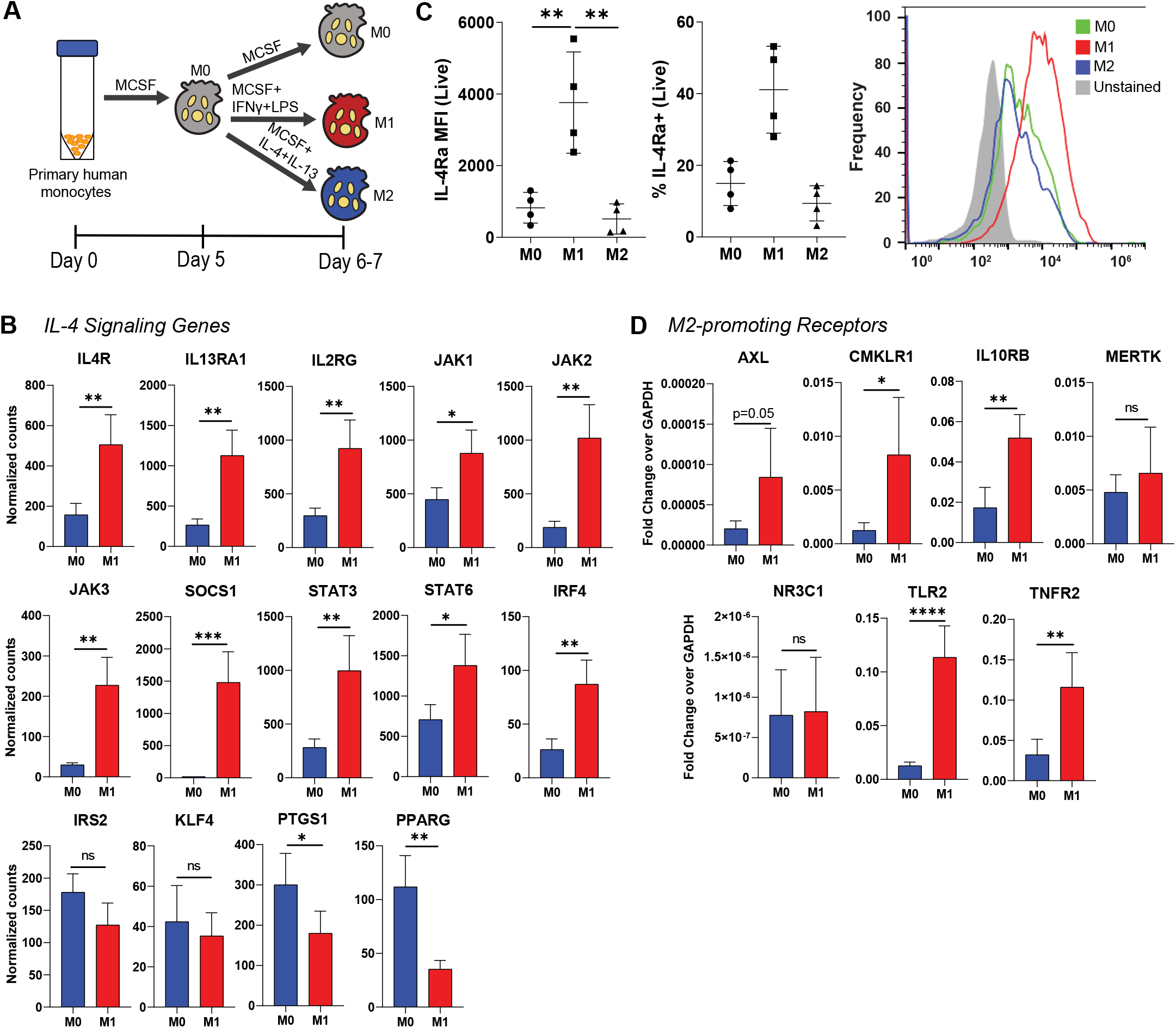
M0 and M1 macrophage expression of M2-promoting receptors. (A) Polarization of unactivated (M0) macrophages to M1 or M2. (B) NanoString counts of genes associated with the IL-4 signaling pathway, using cells collected 24 hours after the addition of polarizing cytokines (day 6). Data are represented as mean ± SD. Unpaired t-test, n=4 donors, *p<0.05, **p<0.01, ***p<0.001. (C) Flow cytometric analysis of IL-4Rα expression, using cells collected at day 7. Data are represented as mean ± SD. One way ANOVA with Tukey’s post-hoc, n=4 donors, **p<0.01. (D) qRT-PCR analysis of receptors associated with promoting an M2-like macrophage phenotype, using cells collected at day 7. Data are represented as mean ± SD. Unpaired t-test, n=3-4 donors, *p<0.05, **p<0.01, ****p<0.0001.

### NanoString gene expression assay

RNA was extracted from cells using the RNAqueous-Micro Total RNA Isolation Kit (ThermoFisher, Waltham, MA, U.S.) and RNA concentration was measured using the Nanodrop ND1000. Gene expression was measured with a custom-designed 72-gene panel (Supplemental Table 1) representing genes associated with typical M1 and M2 phenotype markers as well as angiogenesis and fibrosis (NanoString Technologies, Seattle, WA, U.S.), using 100 ng of RNA per sample, according to NanoString’s protocol. Quality control analysis was conducted and raw data were normalized to in-house positive controls using nSolver 4.0 software (NanoString Technologies, Seattle, WA, U.S.) as recommended by the manufacturer.

### Flow cytometry

Macrophages were incubated with Fc block (BD Biosciences, Franklin Lakes, NJ, U.S.) for 10 minutes at 4°C, then incubated with primary-conjugated antibodies (Biolegend, San Diego, CA, U.S.) and viability stain (Invitrogen, Carlsbad, CA, U.S.) for 15 minutes at 4°C. Antibodies used to stain for IL-4Rα were APC anti-human CD124 (IL-4Rα) (Biolegend, clone G077F6, 1:100) and Live/Dead Fixable Green (Invitrogen, 1:200). To amplify IL-4Rα signal, APC FASER kit (Miltenyi Biotec, Bergisch Gladbach, Germany) was used for two cycles, according to kit instructions. Viability stain used to stain macrophages for phagocytosis was Live/Dead Fixable Aqua (Invitrogen, 1:200). Samples were fixed and permeabilized using the Foxp3/Transcription Factor Staining Buffer Kit (Tonbo Biosciences, San Diego, CA, U.S.), then incubated with primary-conjugated antibodies for 45 minutes at 4°C. PE anti-human CD68 (Biolegend, clone Y1/82A, 1:100) was used to stain macrophages for phagocytosis. Data was measured on the Accuri C6 (BD) or LSR II (BD) flow cytometer and analyzed using FlowJo v10 software (BD).

### Quantitative reverse transcription polymerase chain reaction

RNA was extracted using the RNAqueous-Micro Total RNA Isolation Kit and RNA concentration was measured using the Tecan Infinite M200 microplate reader. cDNA was synthesized using the High-Capacity cDNA Reverse Transcription Kit (Invitrogen). qRT-PCR was conducted using custom oligonucleotides (ThermoFisher, Supplemental Table 2) and SYBR Green Master Mix (ThermoFisher). Statistical testing was performed using log-transformed data.

### Enzyme-linked immunosorbent assays (ELISAs)

Cell-conditioned media was collected 24 hours after the addition of polarizing cytokines and frozen at -80°C until ELISAs were run. ELISAs were conducted using kits from Peprotech (PDGF-BB, CCL5, and VEGFA) and R&D Systems (Minneapolis, MN, U.S.) (CCL17). Absorbance was measured on the Tecan Infinite M200 microplate reader.

### Migration assays

#### Macrophage migration in response to SDF-1α

6.5 mm polyester transwell membranes with 8.0 µm pores (Corning, Corning, NY, U.S.) were coated with 5 µg/cm^2^ of bovine type I collagen (Gibco, ThermoFisher) for one hour and placed in a 24-well plate. Macrophages were suspended in serum-free media at a concentration of 500,000 cells/mL and 100 µL were added to each transwell. After allowing cells to settle for 10 minutes, 600 µL of media supplemented with 10 ng/mL of recombinant human stromal derived factor-1α (SDF-1α) was added to the bottom chambers of the experimental group. 600 µL of basal media was added to the bottom chambers of the negative control group. Samples were incubated for 24 hours, after which media and remaining cells in the upper chamber were removed with sterile cotton swabs, while cells adhered to the lower membrane were fixed with ethanol and stained with 0.2% crystal violet solution (Fisher Chemical, Hampton, NH, U.S.). Images of the transwells were taken on an inverted microscope set to 10x objective and cells were counted using ImageJ software (NIH, Bethesda, MD, U.S.).

#### Endothelial cell migration in response to macrophage-conditioned media

Macrophage-conditioned media without the influence of polarizing cytokines was generated by incubating polarized macrophages in basal media for 24 hours, then frozen at -80°C until assay was conducted. Human umbilical vein endothelial cells (HUVECs) were suspended in serum-free media at a concentration of 500,000 cells/mL and 100 µL were added to each transwell. After allowing cells to settle for 10 minutes, 600 µL of macrophage-conditioned media was added to the bottom chambers. Samples were incubated for 24 hours, after which cells on the bottom of the transwell were stained and counted as described above.

### MTT metabolism assay

HUVECs were seeded in 96-well plates at 50,000 cells/well and cultured in 100 µL of macrophage-conditioned media for 24-48 hours. Media supplemented with 10 ng/mL of recombinant human VEGF (GenScript, Piscataway, NJ, U.S.) was used as a positive control. 10 µL of MTT reagent from the MTT Cell Proliferation Assay Kit (Cayman Chemical, Ann Arbor, MI, U.S.) was added to each well of one plate and incubated for 4 hours. 100 µL of sodium dodecyl sulfate from the same kit was then added to each well and incubated for an additional 18 hours. Absorbance at 570 nm was measured on the Tecan Infinite M200 microplate reader.

### Phagocytosis

Viable HL-60 cells were labelled with CellTrace Far Red (Invitrogen) according to manufacturer instructions, then serum-starved for 2 hours. Hydrogen peroxide (Fisher Chemical) was added to the cell suspension at a concentration of 800 µM for 3 hours to induce apoptosis. Macrophages were co-cultured with either apoptotic HL-60s or crimson FluoSpheres (Invitrogen) for 3 hours to facilitate phagocytosis. Samples were immediately stained for CD68 and Live/Dead for flow cytometry analysis as described above.

### Statistics

Most results were analyzed using GraphPad Prism 8 software (GraphPad Software, San Diego, CA, U.S.). Heat map and dendrogram were generated using R software.

## Results

### M1 activation increases expression of IL-4Rα, IL4 signaling genes, and other M2-promoting receptors

In order to test the potential of M0 and M1 macrophages to switch to the M2 phenotype, we determined their expression of receptors known to promote the M2 phenotype. As IL-4 is the primary cytokine used to polarize M2 macrophages in vitro to model macrophages in late-stage tissue repair, we used NanoString multiplex gene expression analysis to measure the expression of the three components of the IL-4 receptor complexes (IL4R, IL13RA1, IL2RG) as well as their corresponding Janus kinases (JAK1, JAK2, JAK3) and several other genes involved in IL-4 signaling (39). Expression of the receptor complexes, Janus kinases, and other IL-4 signaling genes was significantly upregulated in M1 macrophages compared to M0 macrophages (**Figure 1B**). We then verified surface expression of IL-4 receptor alpha (IL-4Rα) on M1 macrophages via flow cytometry. Mean fluorescence intensity (MFI) of IL-4Rα was significantly higher on M1 macrophages than both M0 and M2 macrophages, as was the percentage of IL-4Rα-positive cells (**Figure 1C**).

In order to determine if this upregulation of M2-promoting genes and proteins is exclusive to the IL-4 pathway, we used qRT-PCR to measure gene expression of an additional repertoire of receptors involved in M2 polarization outside of the IL-4 pathway, including AXL, MERTK, CMKLR1, IL10RB, TLR2, TNFR2, and NR3C1 (31, 40-44). All receptors except MERTK and NR3C1 were upregulated in M1 macrophages compared to M0 (**Figure 1D**). Together, these data indicate that M1 macrophages may be primed to switch to the M2 phenotype via upregulation of M2-promoting receptors, particularly IL-4Rα.

### M1 macrophages are not more sensitive to IL-4 than M0 macrophages

Having confirmed upregulation of IL-4Rα in M1 macrophages, we hypothesized that this increased expression would translate to increased sensitivity to IL-4. To test this hypothesis, we first compared the response of M0 and M1 macrophages to low doses of the cytokine. After 6 hours of exposure to varying doses (10, 1, 0.1, and 0.01 ng/mL), we used NanoString to measure the expression of a panel of genes including M1 markers, IL-4 signaling genes, and the M2 markers CCL22 and CD206. We expected the M1 macrophages to upregulate M2 genes at lower doses compared to the M0 group. In contrast, IL-4-treated M0 macrophages exhibited higher CCL22 expression at the 10 ng/mL dose and higher CD206 expression at doses as low as 0.1 ng/mL (**Figure 2B**). We then tested whether M1 macrophages would upregulate M2 genes more rapidly than M0 macrophages after IL-4 treatment. Using the same gene panel with an additional M2 marker, CCL17, we treated M0 and M1 macrophages with 10 ng/mL of IL-4 and tracked the expression of these genes over the course of 24 hours, expecting M1 macrophages to upregulate M2 marker genes at earlier time points. Instead, we found that the two groups upregulated different M2 markers within 24 hours of IL-4 exposure. While IL-4-treated M0 macrophages exhibited increased expression of CD206 and CCL22 immediately after stimulation, IL-4-treated M1 macrophages upregulated CCL17, which increased for the duration of the study (**Figure 2C**). Additionally, M1 genes were downregulated in IL-4-treated M1 macrophages within 24 hours, confirming a switch in phenotype (**Figure 2D**). All but two of the fourteen IL-4 signaling genes evaluated were expressed at higher levels by M1-derived M2 macrophages for the 24 hour period, although they generally decreased during this time frame (**Figure 2E**). Therefore, the increased expression of M2-promoting receptors did not lead to a more sensitive or faster response to IL-4 treatment, but IL-4 treatment did cause upregulation of different M2 markers in M1 macrophages compared to M0.

**Figure 2.**
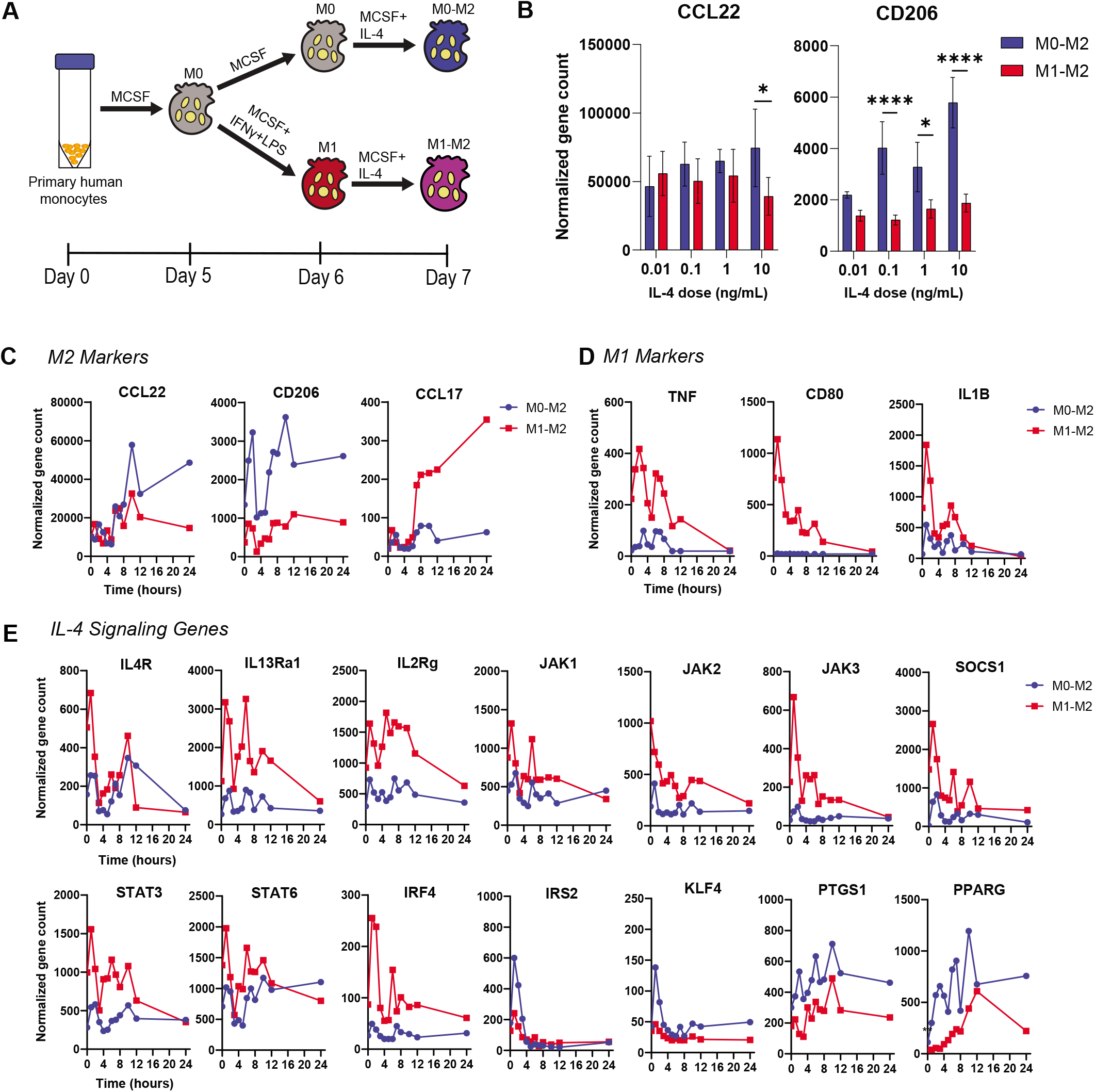
Response of M0 and M1 macrophages to IL-4. (A) Treatment of M0 or M1 macrophages with IL4 to generate M0-derived M2 (M0-M2) or M1-derived M2 (M1-M2). (B) NanoString counts of M2 genes after 6 hours of exposure to varying doses of IL-4. Data are represented as mean ± SD. 2-way ANOVA with Sidak’s test, n=4 donors, *p<0.05, ****p<0.0001. (C) Nanostring counts of genes associated with M2 phenotype, (D) M1 phenotype, and (E) IL-4 signaling pathway over the course of 24 hours of exposure to 10 ng/mL of IL-4. Data are represented as mean only, n=4 donors.

### M0-derived and M1-derived M2 macrophages are distinct phenotypes

These unexpected results led to a new hypothesis that M1 activation changes the subsequent response to IL-4, as measured by differences in expression of M2 markers and possibly other genes. Using an expanded NanoString panel of genes including macrophage phenotype markers, genes associated with angiogenesis, and genes associated with tissue deposition or fibrosis, we evaluated the gene expression profiles of M0 and M1 macrophages treated with 10 ng/mL IL-4 for 24 hours (M0→M2 vs. M1→M2, respectively). M1→M2 macrophages expressed higher levels of several genes compared to M0, M1, and M0→M2 macrophages, including CCL17, CCL5, CXCR4, JAG1, VEGFA, and PDGFB (**Figure 3A**). M0→M2 macrophages, on the other hand, expressed higher levels of CD206, DACT1, and HSPG2. Both the M0→M2 and M1→M2 groups upregulated CCL22 and FLT1 (VEGF receptor-1) compared to the M0 and M1 controls, but were not significantly different from each other. Hierarchical clustering of the entire 72-gene panel data set established that the M1→M2 group, while more closely related to the M0→M2 group than to the M0 or M1 controls, clustered separately from the M0-M2 group and is therefore phenotypically distinct (**Figure 3B**).

**Figure 3.**
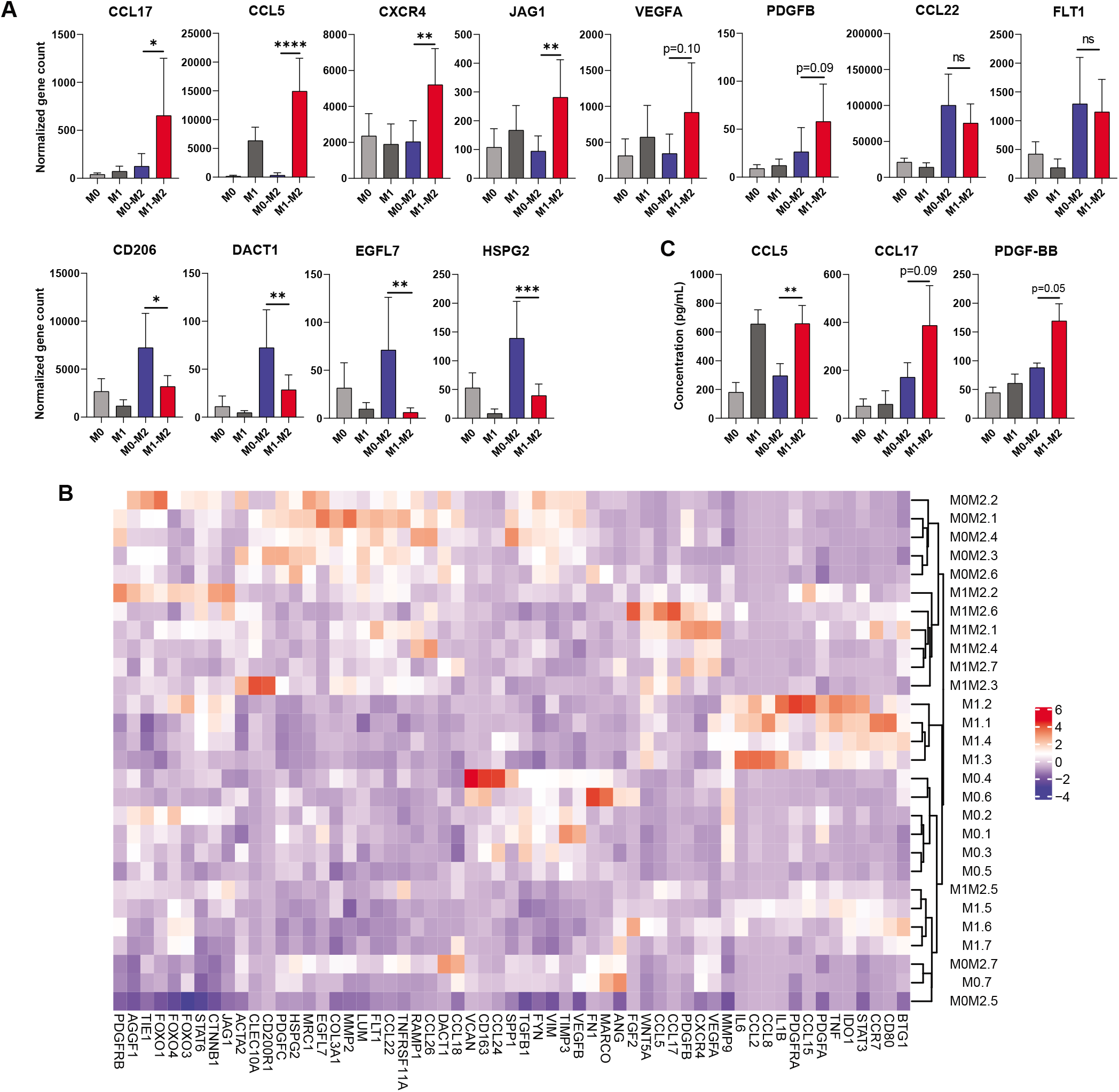
Differential gene expression in M1→M2 macrophages. (A) NanoString gene counts of highly expressed markers associated with M2 phenotype, angiogenesis, and fibrosis. Data are represented as mean ± SD. One-way ANOVA with Tukey’s post-hoc, n=7 donors, *p<0.05, **p<0.01, ***p<0.001, ****p<0.0001. (B) Hierarchical clustering of entire data set. (C) Protein secretion of selected highly-expressed M1→M2 genes. ELISA analysis of macrophage-conditioned media. Data are represented as mean ± SD. One-way ANOVA with Tukey’s post-hoc, n=3-4 donors, ***p<0.001. Only significant differences between M0-M2 and M1-M2 are depicted, although differences were detected between other groups. Figure 3a: M0-M2 and M1-M2 expression of M2, angiogenesis, and tissue deposition genes Statistics: One-way ANOVA with Tukey’s post-hoc; n=7 donors; *: p < 0.05, **p<0.01, ***p<0.001, ****p<0.0001; mean w/SEM

In order to determine whether increased gene expression of CCL17, CCL5, VEGFA, and PDGF-BB translated to increased protein secretion, we conducted ELISAs to measure concentration of these proteins in the culture media after polarization. As expected, M1→M2 macrophages secreted higher levels of CCL17, CCL5, and PDGF-BB compared to M0→M2 (**Figure 3C**). Results of the VEGFA ELISAs, on the other hand, exhibited high variability and no significant differences were found (Supplemental Figure 1).

Collectively, these results indicate that M2 macrophages that were previously M0 or M1 exhibit distinct phenotypes.

### M0→M2 and M1→M2 macrophages exhibit distinct functions

Finally, we set out to evaluate potential differences in functional behavior between M1→M2 and M0→M2 macrophages. Because a critical process in the early inflammatory response is the recruitment of macrophages to injured cells in large part through SDF-1α, and because M1→M2 macrophages upregulated expression of CXCR4, the SDF-1α ligand, compared to M0→M2 macrophages, we first tested the ability of these cells to migrate in response to SDF-1α. Using a transwell migration assay, we measured the migratory response of macrophages to basal media or to media containing SDF-1 α. M0→M2 macrophages were more motile than M1→M2 macrophages, migrating more in basal media (**Figure 4A**). However, the inclusion of SDF-1 α in the media increased transwell migration of M1→M2 macrophages, while it had no effect on M0→M2 macrophages (**Figure 4B**).

**Figure 4.**
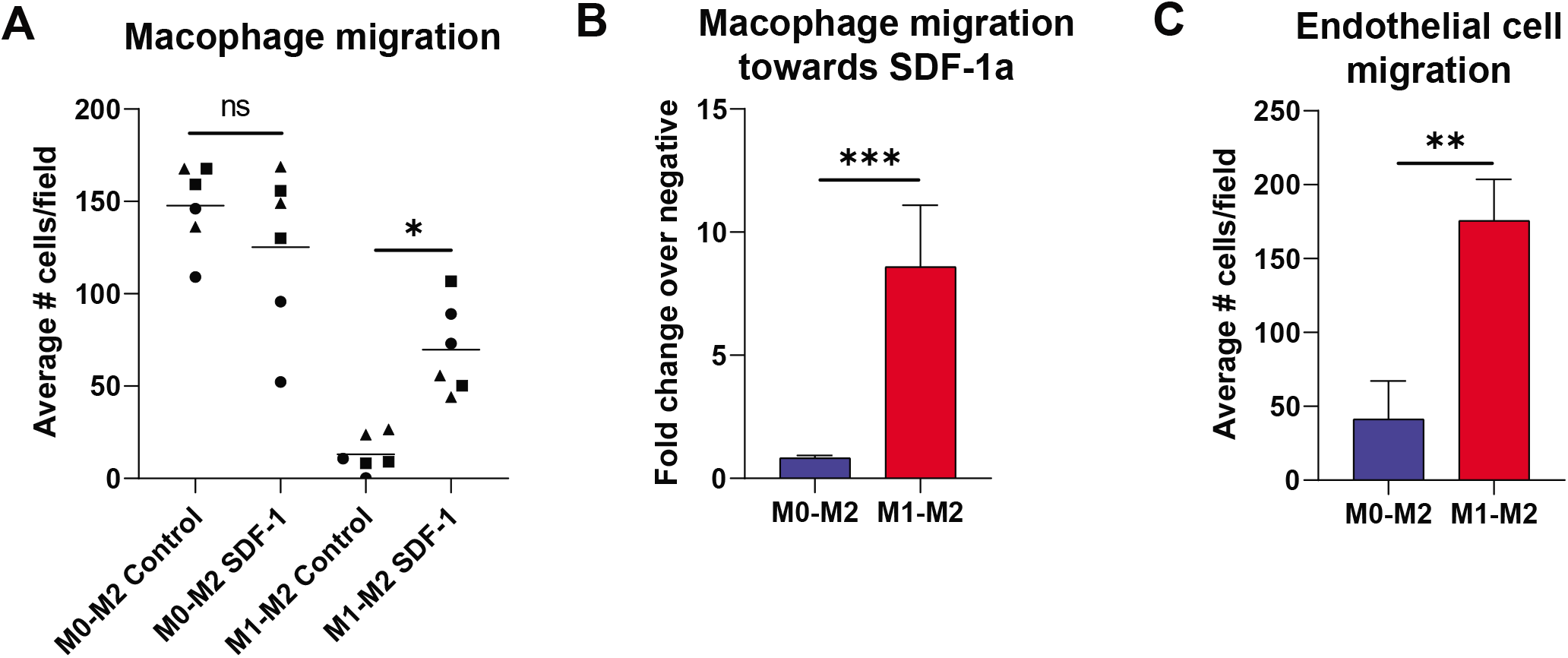
Functional assays in M0→M2 and M1→M2 macrophages. (A) Migratory response of M0→M2 and M1→M2 macrophages towards media ± SDF-1. Data are represented as single replicates with mean only. One-way ANOVA with Tukey’s post-hoc, n=6 replicates from 3 donors depicted with different symbols, *p<0.05. (B) Migratory response of macrophages towards SDF-1, shown as fold change of experimental cell counts over blank negative control cell counts. Data are represented as mean ± SD. Unpaired t-test conducted on log-transformed data, n=6 replicates from n=3 donors, ***p<0.001. (B) Endothelial cell migration towards macrophage-conditioned media. Data are represented as mean ± SD. Unpaired t-test, n=6 replicates from n=2 donors, **p<0.01.

Again using a transwell migration assay, we next found that conditioned media from M1→M2 macrophages recruited more HUVECs compared to M0→M2 macrophage-conditioned media (**Figure 4C**). We also measured proliferation of HUVECs over 24 and 48 hours of incubation in macrophage-conditioned media, but no significant differences were observed between the HUVECs incubated in media from M0→M2 or M1→M2 macrophages (Supplemental Figure 2).

Macrophages are well-known for their role as phagocytes, and some studies have shown increased phagocytosis among M2-activated macrophages (45, 46). In order to assess differences in phagocytic activity between M0→M2 and M1→M2 macrophages, we cultured each group for 3 hours with either fluorescently labelled latex beads or apoptotic HL-60 cells. Using flow cytometry to identify target-containing macrophages, we found no significant differences between the groups in percentage of target-positive cells or MFI of target (Supplemental Figure 3).

## Discussion

We have established here that M1 activation primes macrophages to polarize to an M2-like phenotype that exhibits unique gene and protein signatures compared to M2 macrophages derived from an unactivated state. M1→M2 macrophages were more migratory towards SDF-1 α, suggesting an enhanced ability to traffic towards sites of injury, and induced more HUVEC migration than did M0→M2 macrophages, suggesting that they may play an important role in angiogenesis. The characterization of this unique phenotype points to M1 activation as a major influence on subsequent M2 macrophage behavior. These results have critical implications for understanding how macrophage phenotype is regulated during tissue repair, and suggest that correcting M1 activation may be a therapeutic target for impaired healing situations characterized by deficient M2 activation.

Several studies have previously established a role for M1 macrophages in early angiogenesis (3, 4, 10), and further investigations have suggested that they may also serve to regulate M2 polarization in late-stage healing. For example, M1 macrophages not only have the capacity to switch to the M2 phenotype (1, 5, 6, 8, 9), but stimulation of M1 activity has also been shown to improve subsequent M2 marker expression (47-49). Likewise, studies in which M1 macrophages were inhibited during early stages of healing resulted in diminished M2 polarization later on (6, 10). Correspondent to these findings, several pathologies in which the M2 response is inhibited, such as diabetic wounds, also initially lack robust M1 activity at early stages (11-16). Exactly how M1 activation influences M2 behavior, though, remains poorly understood. M2-like macrophages that were previously pro-inflammatory and those that were derived from an unactivated state are both present in the site of injury during the later stages of healing (8). Our findings shed light on the M1-to-M2 transition by showing that M0→M2 and M1→M2 macrophages are different phenotypes, and perhaps perform distinct yet complementary functions in vivo.

Surprisingly, although M1 macrophages upregulated M2-promoting receptors, they did not exhibit increased responsiveness to IL-4 in terms of sensitivity to lower doses or more rapid responses. Rather, M1 activation seems to change the response to IL-4 by generating a distinct M2-like phenotype with a unique gene signature and set of behaviors. This altered response may stem from a change in the IL-4 signaling pathway upon M1 activation – for example, STAT6 is more highly expressed in M1 macrophages compared to M0 macrophages, while M0 macrophages exhibit higher expression of PPARγ (Figure 1B). Deeper investigation into IL-4 pathway changes may be required to understand the underlying mechanisms of the M1 macrophage response to IL-4.

This study establishes an initial characterization of M1→M2 macrophages, and shows they are phenotypically distinct from M0→M2 macrophages. In line with their emergence during the later stages of tissue repair and angiogenesis, M1→M2 macrophages upregulate PDGF-BB, essential for the stabilization of nascent vasculature, and CCL17, which may promote the resolution of inflammation via regulatory T cell recruitment. Though promising, these findings must be investigated further, potentially via transcriptome sequencing to identify all differences between M0→M2 and M1→M2 macrophages, as well as studies to confirm this phenomenon in vivo.

In order to design therapies that promote angiogenesis and healing via macrophage modulation, a thorough understanding of regulatory mechanisms during normal tissue repair is critical. Although the effects of M1 activation on M2 polarization were previously unknown, we have shown here that the M1→M2 phenotype is distinct and appears to display important functions for wound healing and angiogenesis.

## Acknowledgements

We thank Jessica Eager for technical assistance with heat map generation and hierarchical clustering data analysis. This study was supported by research funding from NHLBI (grant number R01 HL130037).

## Authorship Contributions

E.M.O. performed the experiments and analyzed results; E.M.O. and K.L.S. designed the research and wrote the paper; K.L.S. supervised the research.

## Abbreviations

IFNγ: interferon-gamma
IL-4: interleukin-4
LPS: lipopolysaccharide
MTT: 3-(4,5-dimethylthiazol-2-yl)-2,5-diphenyltetrazolium bromide
PDGF-BB: platelet-derived growth factor BB
SDF-1α: stromal-derived factor-1 alpha
VEGFA: vascular endothelial growth factor A

