## Supplementary figures and images for "Prior Pro-inflammatory Polarization Changes the Macrophage Response to IL-4"

### Supplemental Figures

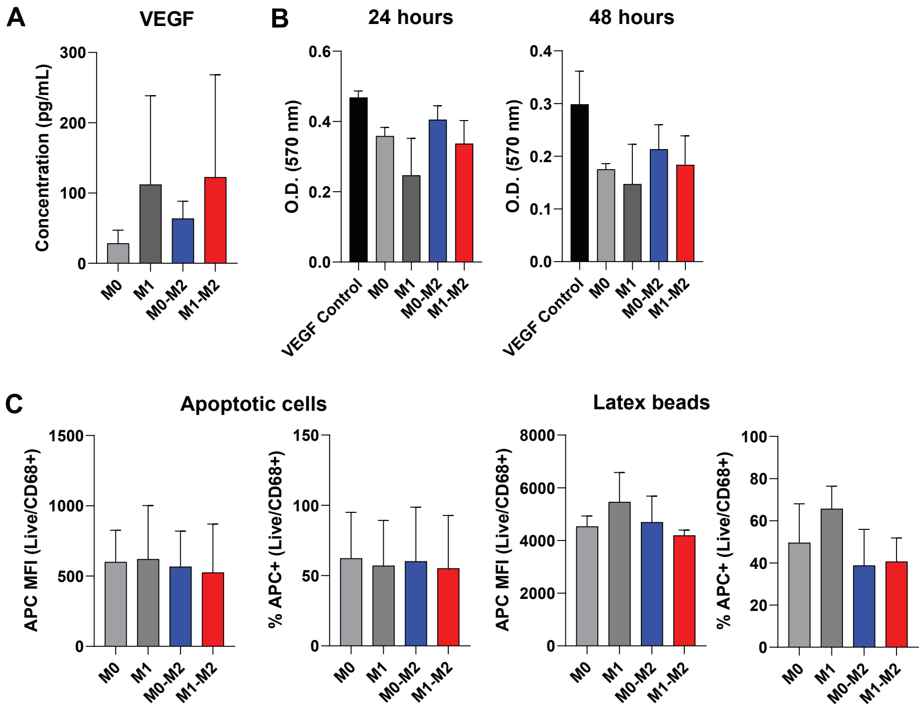
